# Restoration of Susceptibility of *bla*_NDM_-bearing *Escherichia. coli* to Carbapenem Antibiotics by Exogenous *N*-acetylcysteine

**DOI:** 10.1101/2025.03.31.646374

**Authors:** Dexi Li, Yuzhe Zhao, Xinyu Guo, Chenglong Li, Yage Sun, Yixin Wang, Lijuan Chang, Xiaoming Wang, Yanling Wang, Stefan Schwarz, Xiang-Dang Du

## Abstract

The global spread of the New Delhi metallo-β-lactamase (NDM) -producing carbapenem-resistant Enterobacteriaceae pose a serious threat to public health, as NDM and its variants enzymes efficiently hydrolyze almost β-lactam antibiotics. The development of novel antibiotics or antibiotic adjuvants capable of eradicating antibiotic-resistance bacteria through multiple mechanisms represents a promising strategy for reversing antibiotic resistance and preventing the emergence of new resistance. In this study, we utilized high-performance liquid chromatography (HPLC) to demonstrate that *N*-acetylcysteine (NAC) is a prominent reducing metabolite in bacteria harboring NDM-5. Notably, NAC was able to restore carbapenem susceptibility in NDM-producing bacteria in vitro. Subsequently, we further investigated the underlying mechanisms involved. Our findings revealed that NAC exerts its effect through multiple mechanisms that collectively contribute to reversing meropenem resistance. Firstly, NAC may inhibit biofilm formation and reduces polysaccharide production in NDM-positive *Escherichia coli*. and this potential mechanism was verified by transcriptomic analysis, reverse transcription PCR (RT-PCR) and a biofilm restoration experiment involving the addition of citrate cycle intermediate metabolites. Secondly, RT-PCR analysis demonstrated that NAC significantly downregulates the expression of *bla*_NDM-5_. Additionally, NAC exhibited competitive inhibition of NDM activity in vitro. Thirdly, NAC was found to enhance the intracellular accumulation of meropenem, therapy increasing its bactericidal efficacy. These three independent mechanisms offer valuable insights for developing adjuvants. The discovery offers a promising prospect for combating the antibiotic resistance concern. This study paves the way for further exploration and potential clinical applications in the fight against antibiotic-resistant bacterial infections.

## Introduction

The novel Delhi metallo-β-lactamase-1 gene, known as *bla*_NDM-1_ (1), was identified in a carbapenem-resistant *Klebsiella pneumoniae* strain in India. Since then, NDM-1 and its variants, resulting from amino acid substitutions at specific positions, have been identified worldwide. As of April 2023, NDM-1 and thirty-six of its variants have been identified, and their enzymatic properties have been analyzed (2-5). The dissemination of *bla*_NDM-1_ and its variants has been reported to occur via various types of plasmids, transposons and other mobile genetic elements, in more than 60 bacterial species from 11 families (2-5).

NDM-1 and its variants are broad-spectrum carbapenemases with the ability to hydrolyze almost all β-lactams, with the exception of aztreonam. These properties pose the most significant threats to the treatment of infections caused by Gram-negative bacteria. Therefore, there is an urgent need to develop inhibitors or adjuvants to (i) restore the susceptibility of antimicrobial agents, and (ii) prevent the emergence of new antimicrobial resistances.

In general, metabolism regulated by specific metabolic pathways can undergo significant alterations when microorganisms are serially exposed to antibiotics, transitioning from a sensitive to a resistant state. Consequently, the identification of crucial metabolic biomarkers holds great promise for providing novel strategies to control antibiotic resistance. In kanamycin resistant *Edwardsiella tarda* strain (6), there are glucose and alanine suppressed in metabolic profile. As a biomarkers, exogenous glucose and alanine could restore the susceptibility to kancomycin by regulating the TCA cycle. Besides, the others metabolics, fructose (7), pyruvate cycle (8), glycine, serine and threonine (9), glutamine (10, 11), L-alanine (12), and pyruvate (13), promote antibiotic accumulation to restore the susceptibility to multidrug resistant bacteria. Herein, we elucidate the pivotal role of metabolic *N*-Acetylcysteine (NAC) to combat antibiotic resistance in bacteria harboring NDM. Our study focuses on the synergistic interaction between meropenem and NAC and the underling molecular mechanism in treating infections caused by NDM-producing *Escherichia coli*.

## MATERIALS AND METHODS

### Strains, media, and reagents

In total, six strains including NDM-1 (n=2), its variants NDM-5 (n=2) and NDM-9 (n=2) from *E. coli* were used for this study. All these strains were collected by the College of Veterinary Medicine, Henan Agricultural University, Zhengzhou, China and were identified by whole genome sequencing and 16S rRNA sequence. Meropenem and 19 commercially available *N*-acetyl amino acids excluding, threonine were purchased from Shanghai Macklin Biochemical Technology joint stock company (Shanghai, China). BCECF-AM fluorescent probe was purchased from AAT Bioquest Inc (California, USA), propidium iodide fluorescent probe was purchased from Beijing Solarbio Science & Technology Co. Ltd (Beijing, China). NPN fluorescent probe was purchased from Shanghai yuanye Bio-Technology Co. Ltd (Shanghai, China)., ATP detection kit was purchased from Beyotime Biotech Inc (Shanghai, China)., EtBr fluorescent detection probe, CCCP efflux pump reagent were purchased from Med Chem Express LLC (Beijing, China), DiSC_3_(5) fluorescent probe, TRIzon Universal RT-PCR kit was purchased from Jiangsu ComWin Biotech Co, Ltd (Jiangsu, China).

### Antimicrobial susceptibility testing and FIC assay by chequerboard

To evaluate the effect of the meropenem antibiotic and 19 commercially available *N*-acetyl amino acids excluding, threonine, the MIC was tested by the broth microdilution method for the Minimal Inhibitory Concentrations (MIC) (14), and the synergistic activities were evaluated by chequerboard analysis. *E. coli* ATCC 29213 served as quality control. In the FIC assay, the set concentrations of meropenem ranged from 0.25 mg/L to 256 mg/L for *bla*_NDM_-positive strains, and NAC ranged from 1.2 mmol/L to 153.6 mmol/L for all strains, respectively. The evaluation standard of thefractional inhibitory concentration (FIC) index was as follows: ≤ 0.5, synergistic effect; 0.5-4.0, irrelevant effect, ≥ 4.0, antagonistic effect.

### Time-kill experiments

The synergistic activity of NAC against *bla*_NDM_-positive *E.coli* was evaluated through time-kill experiments, incorporating minor modifications from previous research (15). Specifically, the *bla*_NDM_-positive *E.coli* strain 1911034 and *E. coli* 6Z13F were cultured overnight, diluted into fresh Brain Heart infusion (BHI) broth, and then incubated for 24 hrs at 37°C. The experiment tested meropenem (at a concentration of 8 mg/L, suitable for carbapenem-resistant strains) and two concentrations of NAC (2.40 and 4.80 mmol/L) both individually and in combination. Viable cell counts were determined at the start of the experiment and at 0, 4, 8,12, and 24 hrs following exposure to either the single antibiotic or the antibiotic in combination with NAC. Biological replicates were performed in triplicate (n=3).

### Assay for resistance evolution

*E.coli* 1911034 were cultured to the logarithmic phase growth phase, as indicated by an optical density at 600 nm (OD₆₀₀) within the range of (0.6 to 0.8). Subsequently, the bacterial suspension was diluted 1:100 into fresh BHI broth. Two treatment groups were established, including Cultures exposed to 1/8×MIC of MEM and Cultures treated with a combination of 1/8×MIC of MEM and 2.40 mmol/L of NAC. Each treatment group was incubated at 37°C for 24 hrs. Following incubation, 1 mL of each culture was transferred to fresh LB broth containing the same drug concentrations to initiate the next passage. This serial passage process was repeated daily for 30 consecutive days. At both day 0 (initial) and day 30 (final), bacterial suspensions were prepared in triplicate and subjected to a standard broth microdilution assay to determine the MIC of meropenem. The fold increase in MIC was calculated using the formula: Fold increase= MIC_final_ / MIC_initial._

### Antibiofilm assays

The potential synergism of meropenem plus NAC combinations against biofilms was investigated using a previous method with minor modifications (16). Bacteria were collected at the logarithmic growth phase (OD 0.5-0.8), centrifuged at 700×g for 5 min, the cell pellet was washed twice after being dissolved in PBS. After resuspending in LB broth, 100 μL were inoculated into each well of a 96-well plate. These samples were divided into three groups (the control group without antibiotic or NAC, the MEM (8 mg/L) group, and the MEM (8 mg/L) plus NAC (4.80 mmol/L) group) and incubated at 37°C for 48 hrs.

After 48 h, the 96-well plate was rinsed with phosphate-buffered saline (PBS), and the supernatants were discarded. Next, 100 μL 1% crystal violet solution was added to each well and stained for 20 min, then washed with PBS, and 100 μL ethanol absolute was added to each well of the plate to dissolve the crystal violet. The absorbance at OD_570_ was recorded with a multimode microplate reader (Sigma-Aldrich, Darmstadt, Germany).

Polysaccharides is the component consistant of biofilm, change of which was also assayed by the following methods with minor modifications (16): Firstly, bacteria were cultured overnight, and then transferred into 10 ml LB medium at a ratio of 1:1000. Bacteria were incubated at 37°C for 48 hrs, and these sample were divided into three treatment groups including no treatment, MEM (8 mg/L) treatment and MEM (8 mg/L) plus NAC (2.40 mmol/L) treatment. Secondly, the cultured bacteria were centrifuged at 25°C and 5000 *g* for 20 min, the supernatants were discarded and the pellets were resuspended in10 ml PBS (first time), 1 ml NaCl (1.5 M) (second time). Thirdly, the mixtures was transferred to a 1.5 ml Eppendorff tube (abbrevation as EP) Added 40 μl of EPM, 10 μl of phenol (5%), and 50 μl of sulfuric acid to a 96-well plate, mixed well and incubated at 25°C for 10 min, and monitored of its absorbance at OD_550_ with a multimode microplate reader (Sigma-Aldrich, Darmstadt, Germany).

To evaluate the impact of NAC on biofilm formation and polysaccharide modulation, tricarboxylic acid (TCA) cycle intermediates—citrate (1.3 mmol/L), succinate (2.1 mmol/L), fumarate (2.1 mmol/L), and malate (1.9 mmol/L)—were supplemented into two experimental groups: (1) MEM (8 mg/L) alone and (2) MEM (8 mg/L) combined with NAC (2.40 mmol/L). The experimental procedures followed established methodologies described in prior studies (17, 18).

### Transcriptomic Analysis

*E.coli* 1911034 was inoculated into fresh LB broth and divided into two groups: MEM (8 mg/L) group and MEM (8 mg/L) plus NAC (2.40 mmol/L) group. Then, bacteria that were cultivated for 48 h were collected by centrifugation, washed twice with PBS, lysed in liquid nitrogen and met the requirements of approximate 0.5 g of bacteria for each sample. Then, these samples were sent to the Novogene Corporation Inc. for transcriptomic sequencing and analysis. Each group had three biological replicates.

The evaluation of the differentially expressed genes were carried out using DESeq2 software, with the parameters log2 (Fold Change) ≥ 0 and padj ≤ 0.05.

### RT-PCR analysis

Overninght grown bacteria were transferred into 300 ml fresh LB broth supplemented with the MEM (8 mg/L), and the MEM (8 mg/L) plus NAC (2.40 mmol/L) group at 37℃with shaking at 180 rpm for 48 hrs (according to the set for transcriptome). The bacteria were collected and extracted for total RNA according to the Manufacturer’s Instructions.

Based on transcriptomic analysis, target genes and gene amplification primers were listed in Table S1, Table S2, whereas the 16s rRNA and *dxs* encoding 1-deoxy-D-xylulose-5-phosphate synthase served as the control. The amplificarion of RT-PCR was conducted by Bio-Rad CFX96 Real-Time PCR System (California, USA). The x-fold changes of each gene were analyzed by Bio-Rad CFX Maestro V2.2 software. Real-time RT-PCR assays were applied for quantitative detection of biofilm-associated genes and others DEGs.

### Potential Mechanism Analysis by Measurement the Inhibition Activity of Enzyme

#### NDM-5 enzyme kinetics parameters

To establish an in vitro inhibition evaluation system for NDM-5, this study successfully prepared high-purity NDM-5 protein using a recombinant expression system. Specific primers with restriction endonuclease site for *bla*_NDM-5_ (*Bam*HI-NDM Forward:CGGGATCCGAAAACCTGTATTTCCAAGGCCAGCAAATGGAAACTGGCGAC; *Xho*I-NDM-Reverse: CCGCTCGAGTCAGCGCAGCTTGTCGGCCATG) were designed to clone the *bla*_NDM-5_ gene into the pET28a(+) vector, which was then transformed into *E.coli* BL21(DE3) to construct a His-tag fusion expression system. Further, optimizated the expression condition ion under induction conditions of 37℃, 0.5 mM IPTG for 4 hrs. The target protein was predominantly detected in the supernatant, indicating soluble expression. Protein Purification: Nickel affinity chromatography with gradient elution (20–250 mM imidazole) identified 80 mM imidazole as the optimal elution concentration.

The purified NDM-5 was diluted to 0.05 μM and NAC to 4.8 mmol/L for the following research. A total of 50 μL each of NDM-5 and NAC was added to a 96-well plate, mixed and incubated at 30 ℃ for 20 min. Then, a serial dilution of MEM (final concentrations of 10, 20, 40, 50, 60, 80, 100, 120, 160, 180, 200, and 300 μM) were added to the abovementioned 96-well plate, with three replicates for each group. Its absorbance was detected at OD_300_ by a multimode microplate reader (Sigma-Aldrich, Darmstadt, Germany).

#### The impact of NAC addition to the meropenem-enzyme system

The combination of the NDM-5 (0.05 μM) and NAC (4.80 mM) in a volume of xy µl were incubated at 30℃ for 15 min. Then, 125 μL the mixture of combination and 100 μL MEM (600 μM) were added into a 96-well plate, the absorbance at OD_300_ was monitored with a multimode microplate reader (Sigma-Aldrich, Darmstadt, Germany) for 30 min, and meropenem-enzyme system without NAC was used as the control.

#### IC_50_

A total of 50 μL of NAC solutions with a series of concentrations (19.2, 9.6, 4.8, 2.4, 1.2, and 0.6 mmol/L) was added to a common mixture system containing 50 μL of NDM-5 (200 nM) and 100 μL of HEPES buffer. The mixture was incubated at 37°C for 15 minutes. Subsequently, 100 μL of MEM (400 μM) was added to the mixture, and the absorbance at 300 nm (OD_300_) was measured using a multimode microplate reader (Sigma-Aldrich, Darmstadt, Germany). For controls, 100 μL of NDM-5 was reserved, and 100 μL of MEM (400 μM) was added to 100 μL of HEPES buffer to serve as the negative and positive control groups, respectively.

#### Ki

Based on the data and Lineweaver-Burk double reciprocal plot, the enzyme inhibition kinetics curve was plotted and the K_i_ value was calculated. First, gradient dilutions of MEM (n=6) with an initial concentration of 800 μM were used for the study. Then, 50 μL of NAC (38.4 mmol/L or 19.2 mmol/L), 50 μL of NDM-5 (1 μM) and 100 μL of the pre-diluted MEM concentrations (800 μM to 25 μM) were added to the mixture system in 96-well plates, and the variations of absorbance at OD_300_ were detected by a multimode microplate reader (Sigma-Aldrich, Darmstadt, Germany).

### Assay for Outer membrane permeability, cell membrane integrity, membrane depolarization, proton motive force, and detection for ATP

The assays for outer membrane permeability, cell membrane integrity, membrane depolarization, proton motive force, and detection for ATP were carried out according as described earlier (15).

Bacteria were collected from five different treatment group at the logarithmic growth phase (OD 0.5-0.8), centrifuged at 700×g for 5 min, the cell precipitates were collected and washed twice after being dissolved in PBS, then the optical density at 600 nm (OD_600_) was adjusted to 0.5. These groups included the control group, MEM (8 μg/mL) plus four concentrations (1.2 mmol/L, 2.4 mmol/L 4.8 mmol/L and 9.6 mmol/L) NAC groups.

Fluorescent probe *N*-phenyl-1-naphthylamine, propidium iodide, DiSC_3_(5) and BCECF AM were diluted to a final concentration of 10 μM, 15 μM, 0.5 μM, and 10 μM, respectively, and incubated at 37 °C for 30 min. Then, 200 μL was transferred from each group (with three replicates per group) to a 96-well black reaction plate, and the fluorescence intensity was measured using a multimode microplate reader (excitation wavelength and emission wavelength according to the previous methods) (15). For ATP detection, the variation of total ATP was conducted using the ATP detection kit (Beyotime, Shanghai, China).

### Antibiotic accumulation assay by HPLC

The accumulation of meropenem in *E.coli* 1911034 was detected according to a previously described method (15) with the minor revision about parameters as follows: the concentration of single meropenem group and meropenem plus NAC combination group were 8 μg/mL, 8 μg/mL plus 1.2 mmol/L, 8 μg/mL plus 2.4 mmol/L, and 8 μg/mL plus 4.8 mmol/L respectively. Spursil ODS C18 with mobile phase of methanol/ acetonitrile ethanenitrile/ potassium phosphate monobasic (0.05 M, pH=5.0) (27.2:6.8:166, v/v/v) and the flow rate of 1 mL/min with column temperature of 30 ℃, and the wave length of 300 nm.

## RESULTS AND DISCUSSION

### Enhancing the *In vitro* efficacy of meropenem and preventing resistance evolution against NDM-producing strains enhanced by NAC

There were certain metabolics of *N*-acetyl-amino acids, such as *N*-acetyl-L-aspartic acid confirmed by HPLC-MS/MS in ours previous untargeted metabolomics data (data not shown). To our interesting, some amino acid derivatives can restore susceptibility to meropenem by checkerboard experiments. Therefore, we hypothesis that the combination of *N*-acetyl amino acids with meropenem may result in the eradication of NDM-producing bacteria. Therefore, 19 commercially available *N*-acetyl amino acids excluding, threonine were purchased and subjected to checkerboard analysis. The results revealed that NAC could reverse the meropenem resistance with a minimum Fractional inhibitory concentration (FIC) index in *bla*_NDM-5_-carrying *E.coli* strains 1911034. Furthermore, comparative metabolic profiling of *bla*_NDM-5_-carrying *E.coli* and their *bla*_NDM-5_-negative strains, based on previous published data, consistently revealed that NAC were among the most significantly differentially abundant metabolites (19).

The MIC of meropenem ranged from 16 to 64 mg/L among NDM-positive strains, while the MIC of NAC ranged from 9.6 to 19.2 mmol/L. The FIC demonstrated a significant synergistic effect (FIC ranged from 0.0936 to 0.375) of meropenem in combination with NAC against six carbapenem-resistant strains (**Fig.1**). Bacterial cultures without antibiotic exposure were established as control.

**Fig 1.**
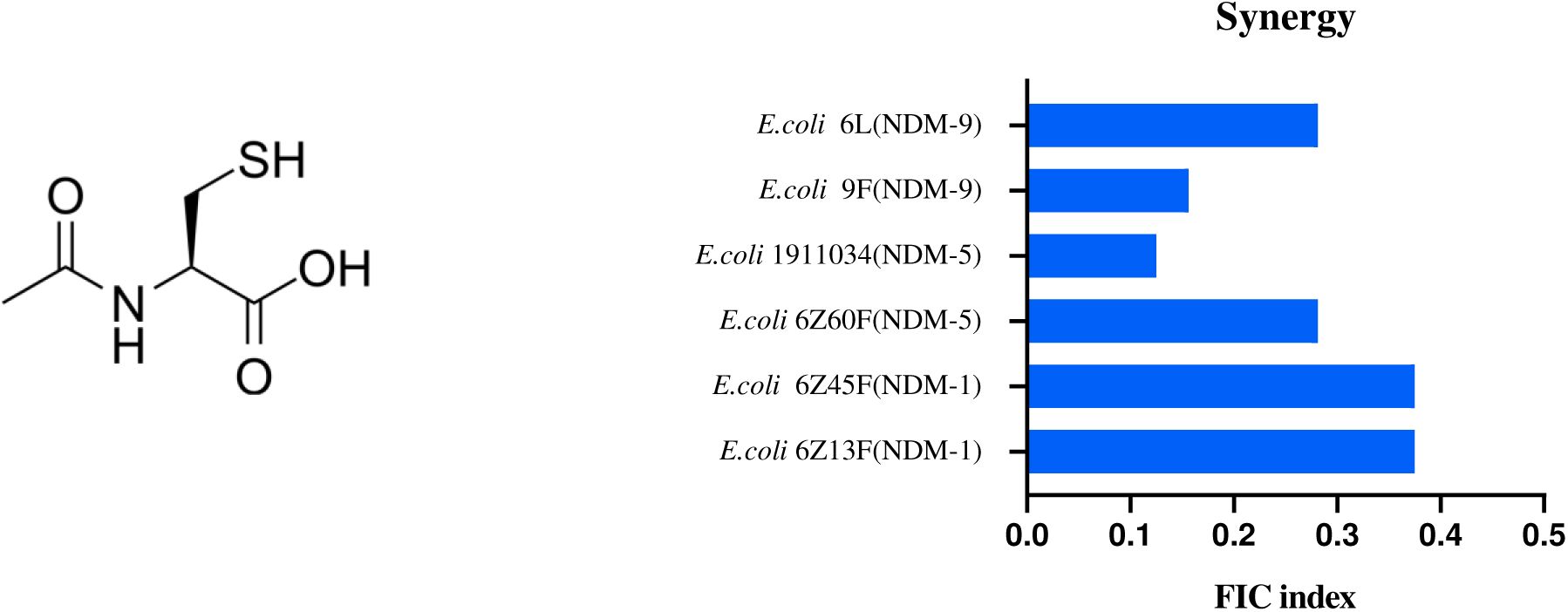
NAC restores the susceptibility of meropenem combating *bla*_NDM_ -bearing bacteria. NAC is a cysteine *N*-terminal acetylation derivative. The FIC assay by Chequerboard were calculated at one quarter of MIC about NAC for different subtyoe *bla*_NDM_ bearing *E.coli*.

NAC, a cysteine *N*-terminal acetylation derivative, is a medication widely used as mucolytic (20). It also possesses antioxidant activity for the treatment of cellular redox imbalance, and anti-inflammatory activity (21). In previous research, NAC demonstrates a dual modulatory role in antimicrobial therapy, exhibiting both synergistic and antagonistic effects (20-27). For Gram-positive bacteria, the combination of NAC with cephalosporins, levofloxacin, showed enhanced activity against antimicrobial resistant *S. pneumoniae* or *S. aureus*, respectively (25, 27). For Gram-negative bacteria, the combination of NAC with CAZ-AVI, meropenem, fosfomycin, or colistin, respectively, could effectively inhibit the growth of resistant strains (20, 22-24, 26, 27).

To reconfirm the effect, the two *bla*_NDM-5_-carrying *E.coli* strains 1911034 and 6Z13F were subjected to growth in the presence of neither meropenem nor NAC (= control group), meropenem (8 mg/L) alone, NAC (either 2.4 or 4.8 mmol/L) alone, and meropenem together with NAC (either 8 mg/L plus 2.4 mmol/L or 8 mg/L plus 4.8 mmol/L). Growth was evaluated after 4, 8, 12, 16, 20 and 24 hrs of incubation. Our time-kill curve assay revealed that the combination of meropenem and NAC exhibited potent synergistic bactericidal activity against *E.coli* 1911034 and *E.coli* 6Z13F (**Fig. 2A** and **Fig. 2B**). Moreover, growth assay in tubes confirmed that there was no visible bacteria growth in meropenem and NAC (8 mg/L and 4.8 mmol/L) combination group (**Fig. 2C** and **Fig. 2D**).

**Fig 2.**
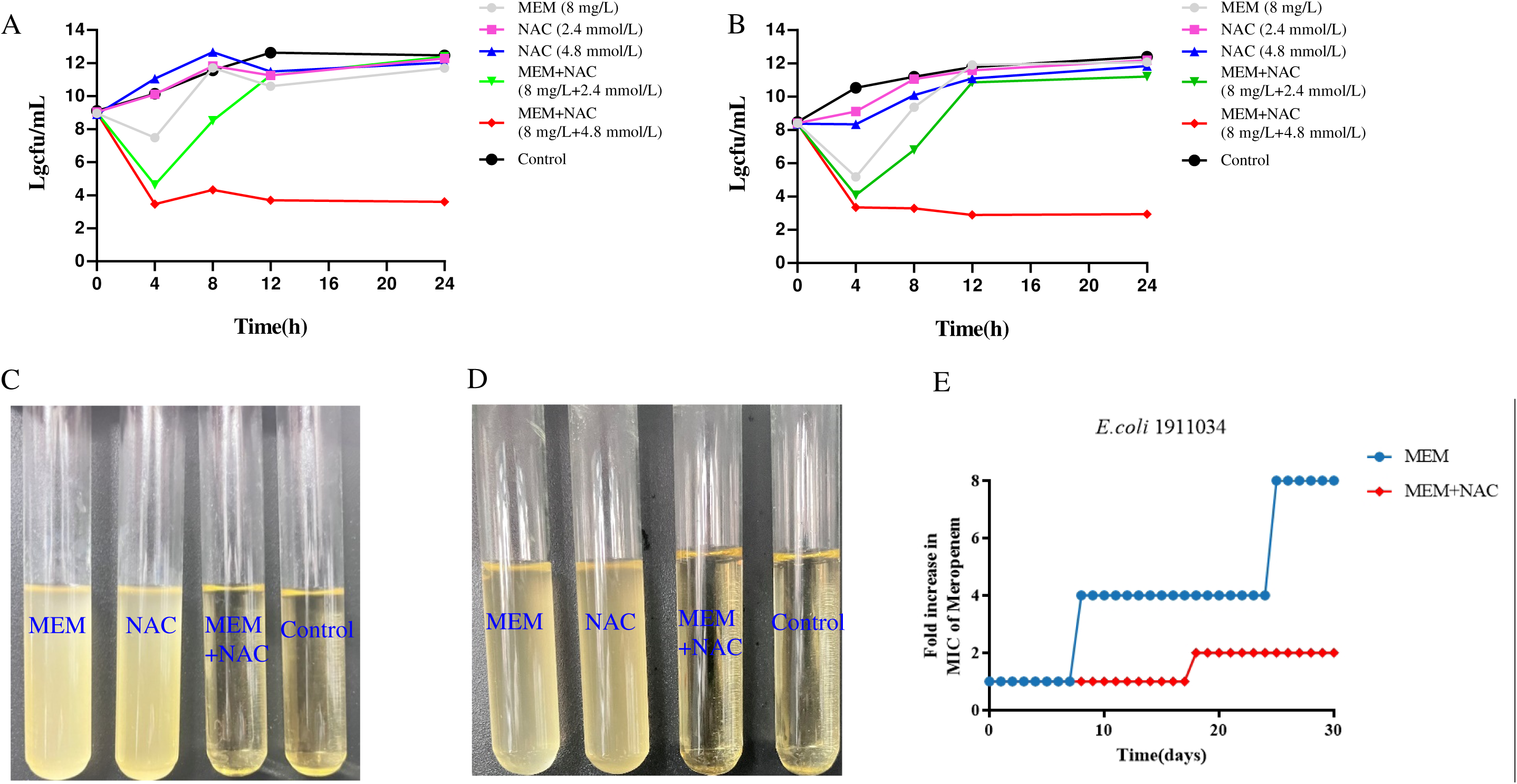
Evaluation of synergistic efficacy by time-killing curve of meropenem-NAC combinations in *E. coli* 1911034 (A) and *E. coli* 6Z13F (B). Both strains were grown to expontial phase and then treated with one meropenem concentration (8 mg/L), two NAC concentrations (2.40 and 4.80 mmol/L) alone or in combination. The combination of meropenem (8 mg/L) and NAC (4.80 mmol/L) had the strongest effect on both strains and killed the bacteria as shown in the time-kill curves (A) and (B) as well as by the lack of visible growth of bacteria in the respective tubes (C) and (D).The potential resistance development was evaluated in *E.coli*1911034 using serial passage experiments. Two treatment groups were compared:(1)a combination group treated with meropenem at 1/8 MIC and NAC at 2.4 mmol/L(MEM+NAC),and(2)a meropenem monotherapy group at 1/8 MIC(MEM) (E).

To evaluate the potential for resistance development, we conducted serial passage experiments over a 30-day period to combination therapy with meropenem at 1/8 MIC and NAC at 2.4 mmol/L, and meropenem monotherapy group at 1/8 MIC. Throughout the experiment, the combination group maintained a relatively stable MIC, with a 2-fold increase in MIC observed by the 18th passage. Whereas, the meropenem single group exhibited a significant 8-fold increase in MIC values (**Fig. 2E**). These findings collectively suggest that the meropenem-NAC combination may effectively kill the NDM bearing strain. This finding provided support for the initial hypothesis, and served as a basis for subsequent investigations into the underlying mechanisms.

### Inhibiton of Biofilm Formation by NAC

Biofilm development is a complex process involving attachment, growth, maturation, and detachment stages. The formation of biofilm not only causes severe problems in bacterial infections, but also results in resistance to some important antimicrobial agents (28). A previous study investigated the molecular mechanisms of synergistic effects of β-lactam antibiotics including ceftazidime-avibactam for ceftazidime-avibactam–resistant enterobacterales, cefditoren for multidrug-resistant *Streptococcus pneumoniae*, ampicillin/sulbactam and meropenem for carbapenem-resistant *K.pneumoniae* or carbapenem-resistant *A.baumannii*, fosfomycin, fluoroquinolones, colistin and other antimicrobial agents in combination with NAC, which revealed that each of these antimicrobial agents in combination with NAC inhibited biofilm formation (16, 20-27, 29). Therefore, biofilm was a crucial target for NAC in enhancing antimicrobial susceptibility or acting as an adjuvant to restore antimicrobial susceptibility.

Based on the results of previous studies, we also hypothesised that NAC might affect the biofilm formation of *bla*_NDM-5_-carrying *E.coli*. In this study, biofilm formation and polysaccharide production change assays were evaluated after different group treatment. In this study, the biofilm production and polysaccharide production analysis were performed using OD-based quantification, following a modified protocol from previous studies (30). Compared to the control group and the meropenem single treatment group, the combination group of MEM plus NAC (= 8 mg/L meropenem plus 4.8 mmol/L NAC) significantly inhibited bacterial growth and biofilm synthesis, and the biofilm reduction ranged from 61.3% to 63.6% (**Fig. 3A**). The polysaccharide production is one of the key components in biofilm formation. In the combination group of MEM plus NAC group, less than 0.085 mg/mL polysaccharide was produced, compared with 0.12 mg/mL in the untreated control group (**Fig. 3B**). Thus, the data obtained from the MEM plus NAC group significantly reduced biofilm formation and polysaccharide production. Citrate-stimulated biofilm formation may be independent of the *ica* locus, and the TCA cycle significantly increases intercellular polysaccharide production in *Staphylococcus epidermidis* (18). We hypothesizes that citrate and others components of the TCA cycle may influence the biofilm formation and the production of polysaccharides. To test this hypothesis, we added citrate, succinate, malate, and fumarate into the MEM group and the MEM plus NAC group. Compared with the MEM plus NAC group, significant increases in biofilm production were observed, ranging from 13.0% to 21.0% at 48 hrs (**Fig. 3C**). Moreover, there was no significant difference in the production of polysaccharides in each group of MEM plus NAC group plus citrate (1.3 mmol/L), succinate (2.1 mmol/L), fumarate (2.1 mmol/L), and malate (1.9 mmol/L), compared with that in the MEM plus NAC group (**Fig. 3D**).

**Fig 3.**
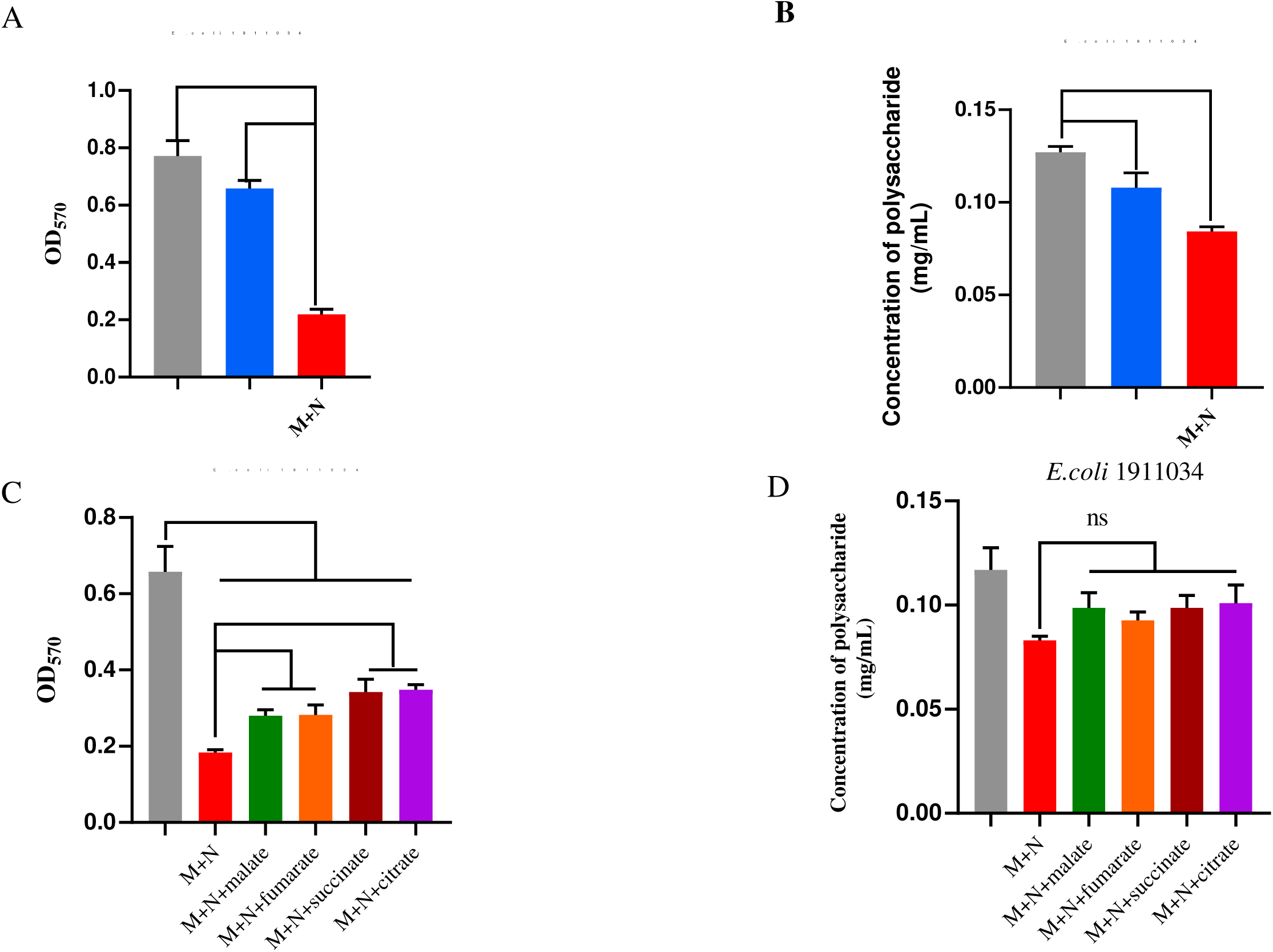
Suppression of biofilm formation by the meropenem-NAC combination (8 mg/L + 4.8 mmol/L). Influence of antibioûlm activity (A) and biofilm change (C). Inf and polysaccharides change (B, D) of meropenem (MEM)/N-acetylcysteine (NAC) combinations alone or together with TCA cycle intermediates citrate (1.3 mmol/L), succinate (2.1 mmol/L), fumarate (2.1 mmol/L), and malate (1.9 mmol/L) against *E. coli* 1911034, respectively. All data obtain from three biology replicates and the significances were evaluated by Prism 8 (GraphPad, USA) (**** P < 0.0001, *** P < 0.001 and ** P < 0.01). M: meropenem; M+N: meropenem plus NAC.

To further examine the potential mechanism of NAC affecting the expression of genes involved in the biofilm formation, transcriptomic analysis of *E.coli* 1911034 was performed after exposure to MEM or to MEM plus NAC for 48 hrs. Compared with MEM single treatment, a total of 807 genes were found to be significantly differentially expressed in the presence of MEM plus NAC (**Fig. 4A**). The horizontal axis represents the log_2_FoldChange value, while the vertical axis represents the-log_10_ (padj) or log_10_ (p-value). 1.301 on the vertical axis is the threshold value, above which genes are considered as up- or downregulated. Among them, 365 genes were up-regulated and 442 genes down-regulated. Gene Ontology (GO) analysis of the up-regulated differentially expressed genes (DEGs) revealed their association with biological processes (e.g., translation, cellular protein metabolic process), cellular components (e.g., ribosome, ribonucleoprotein complex, protein-containing complex), and molecular functions (e.g., structural constituent of ribosome, structural molecule activity) (**Fig. 4B**). Conversely, the down-regulated DEGs exhibited correlations with biological processes (e.g., regulation of RNA metabolic process, cellular process, RNA biosynthetic process), and molecular functions (e.g., oxidoreductase activity). Further, Kyoto Encyclopedia of Genes and Genomes (KEEG) enrichment analysis indicated that significantly up-regulated DEGs were predominantly associated with the ribosome (**Fig. 4C**, **Table S1**, primers listed in **Table S3-1**). Among the 17 pathways to which down-regulated DEGs were associated, eight were correlated with biosynthesis of amino acids and amino acid metabolism (**Table S2**), whereas another three were correlated with tricarboxylic acid (TCA) cycle, fatty acid metabolism and fatty acid degradation (**Fig. 4D**, **Fig 4E**). Within the TCA cycle, the enzymes malate dehydrogenase, citrate synthase, succinyl-CoA synthetase, and succinate dehydrogenase have been observed to reduce the production of succinate, fumarate, malate, oxaloacetate and isocitrate. The transcriptional sequencing analysis revealed that NAC supplementation induces significant downregulation of genes encoding enzymes involved in TCA cycle, which plays a critical role in mediating the observed reduction in biofilm formation. Subsequent experimental validation demonstrated that reintroducing TCA cycle intermediates into the MEM plus NAC group effectively restored biofilm biomass. Gene expression of TCA cycle was modulated by small RNA or the specific site (31-33). These findings collectively confirm our working hypothesis that metabolic impairment of the TCA pathway through NAC exposure represents a key regulatory mechanism underlying the attenuation of bacterial biofilm development. This reduction may, in turn, influence the biofilm formation in *Staphylococcus aureus* isolates (17) and others bacteria including NDM-bearing *E.coli* in this study.

**Fig 4.**
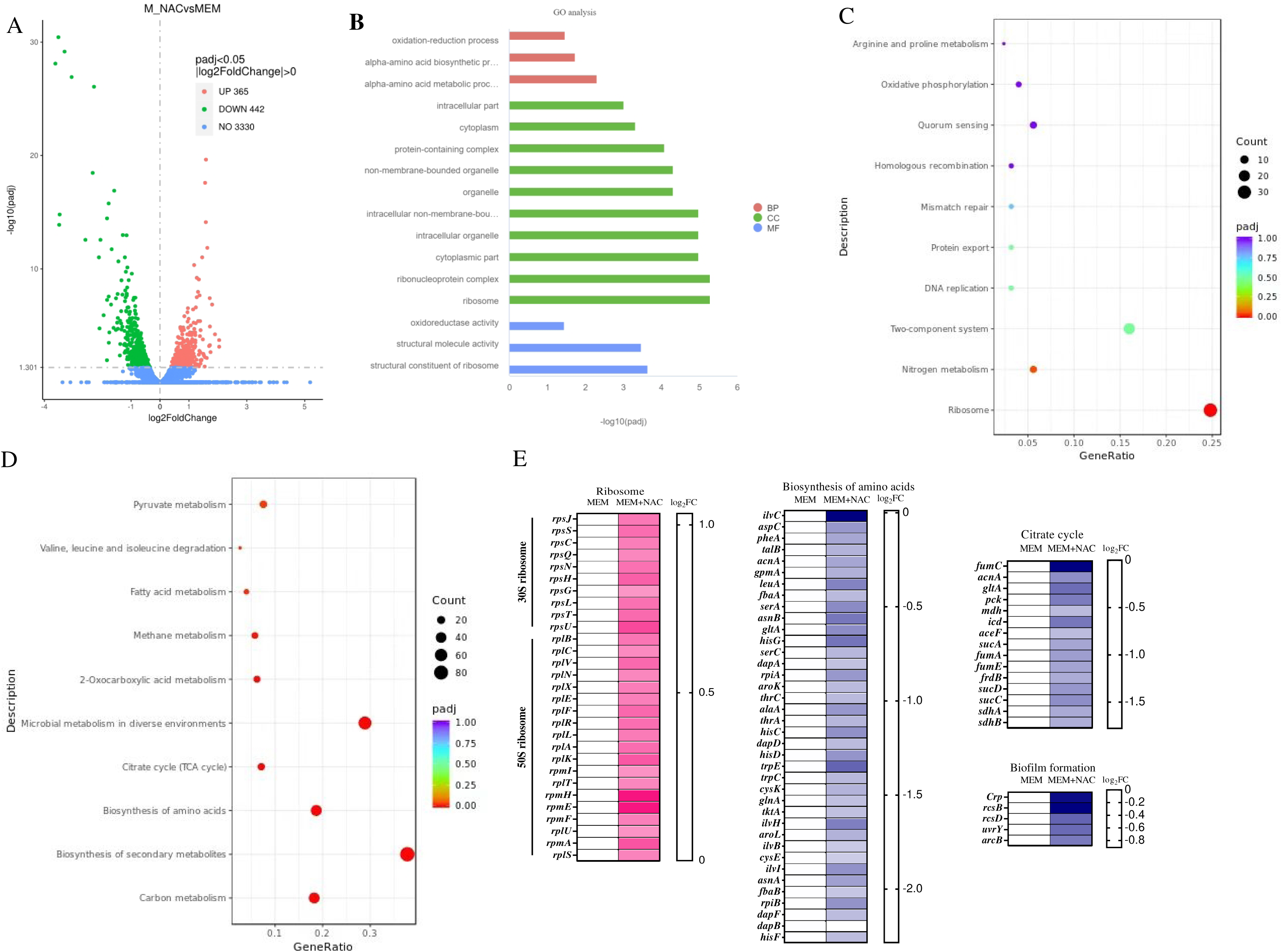
Transcriptome result of *E. coli* 1911034 after treatment with meropenem alone or meropenem couple with.NAC. (A), Up and down change in volcano plot among meropenem alone or meropenem couple with NAC. (B) Gene Ontology (GO) analysis of the differential expression genes (DEGs) reveals their association with biological processes (BP), cellular components (CC) and molecular functions (MF). (C) Upregulation of significant DEGs of Kyoto Encyclopedia of Genes and Genomes (KEEG) enrichment analysis (D) Downregulation of DEG of KEEG, (E) Significant the DEGs include ribosome, biosynthesis of amino acid, citrate cycle and biofilm.

In addition, some of the down-regulated genes are categorized to biofilm pathway-associated *crp*, encoding DNA-binding transcriptional dual regulator CRP, *uvrY* for transcriptional activator UvrY, *acrB* gene for sensor histidine kinase ArcB, *rcsB* for DNA-binding transcriptional activator RcsB, and *rcsD* for RcsD phosphotransferase were confirmed by reverse transcription-PCR (RT-PCR, the primers were listed in **Table S3-2**) (**Fig. 4E**). It is noteworthy that the down-regulated DEGs associated with biofilm regulation exhibited a distribution that was concomitant with the GO pathway of biological processes, cellular components and molecular functions. The down-regulation of energy formation, especially the TCA cycle, may affect the function of anabolism and catabolism, further restricting the function of numerous bacterial proteins, finally resulting in bacterial death.

### Inhibition of *bla*_NDM-5_ gene expression and activity of NDM by NAC

To elucidate the mechanisms by which NAC restores the meropenem susceptibility in NDM-producing bacteria, we hypothesized that NAC may affect the the transcription level of mRNA gene about *bla*_NDM-5_ and or NDM activity. To verify this hypothesis, we evaluated the effect of *bla*_NDM-5_ expression in the mRNA phase and the NDM enzyme itself in vitro after MEM or MEM (8 mg/L) plus lower concentration of NAC (2.40 mmol/L) treatment. In order to obtain the true and relatively reliable result of the variance about *bla*_NDM-5_, two control gene primer pairs for the 16s rRNA gene (**Table S3-2**) and the *dxs* gene, encoding 1-deoxy-D-xylulose-5-phosphate synthase, as well as for the *bla*_NDM-5_ gene were designed for this study (**Table S3-3**). To our surprise, although the relative expression of control genes showed some variations, there were a significant influence of MEM plus NAC (2.40 mmol/L) exposure for 48 hrs on the mRNA levels of the *bla*_NDM-5_-bearing *E. coli* 1911034 strain (**Fig. 5A** and **Fig. 5B**). In previous studies, Metformin (15) and SLAY-P1 (34) have been demonstrated to inhibit the expression of *tet*(A) gene and *vanA* resistance cluster, respectively. This inhibition restores the susceptibility to tetracycline and vancomycin. Similarly, it is hypothesized that NAC may inhibit the expression gene of *bla*_NDM-5,_ thereapy disrupting the synthesis of the NDM-5 protein and potentially reversing carbapenem resistance.

**Fig. 5.**
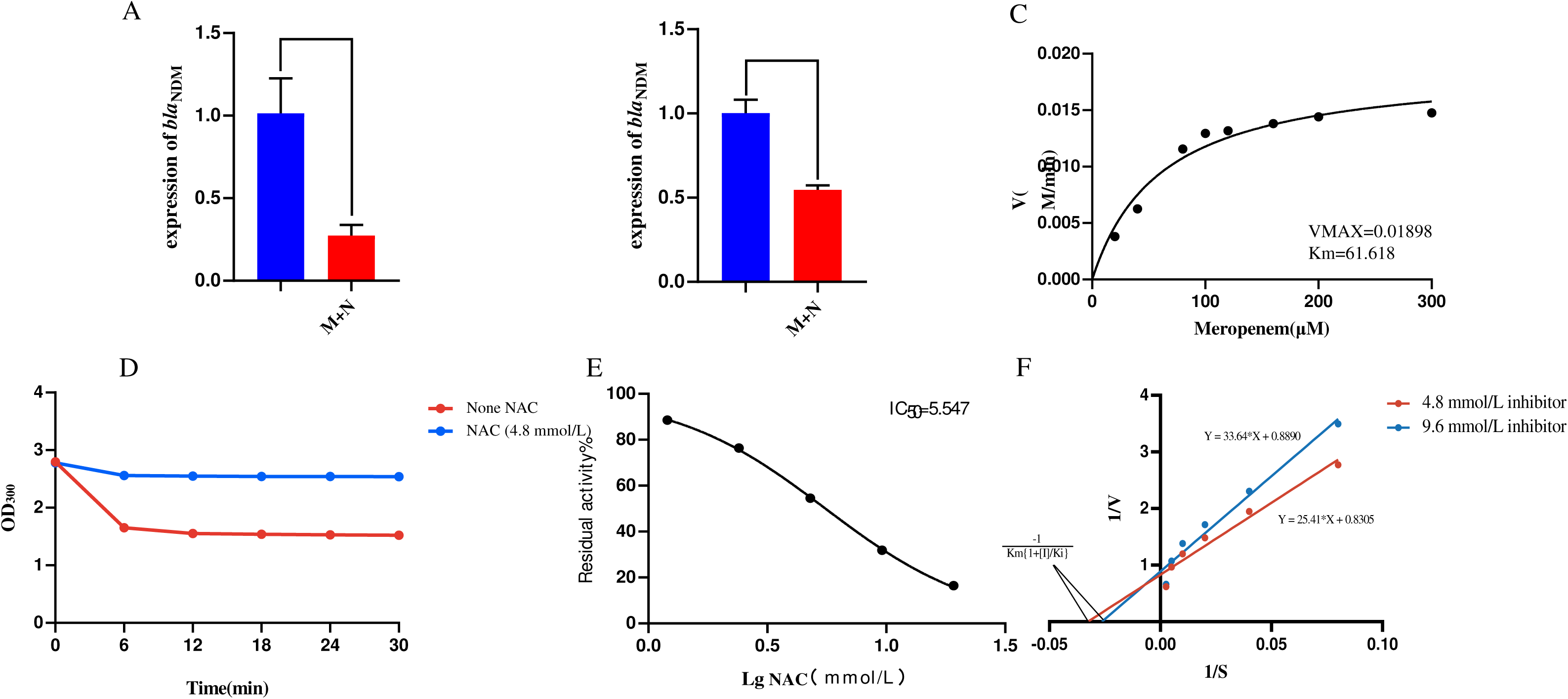
Inhibition of the *bla*_NDM-5_ gene expression and of the activity of NDM-5 by NAC. (A) and (B) represent the expression variation of *bla*_NDM-5_ in comparison to different control genes: 16s rRNA gene (A), *dxs* gene (B); (C) Assay of NDM-5 enzyme kinetics parameters based on the curve of meropenem concentration and NDM-5 enzyme reaction. (D) The impact of NAC addition to the meropenem-NDM-5 enzyme system. (E) Evaluation of IC_50_. (F) The data and Lineweaver-Burk double reciprocal plot were used to set the enzyme inhibition kinetics curve and calculate the Ki value. All data obtain from three biology replicates and the significances were evaluated by Prism 8 (GraphPad, USA) (**** P < 0.0001, *** P < 0.001 and ** P < 0.01). M: meropenem; M+N: meropenem plus NAC.

Then, we explored the inhibition enzyme assays following exposure of NAC interaction with the NDM-5 enzyme. The *bla*_NDM-5_ gene was cloned, expressed and purified using an expression vector BL21 (DE3). As presented in **Fig. 5C**, in comparison with the untreated group, the concentration of meropenem decreased rapidly within six min following the addition of NDM-5 enzymes. The evaluation of inhibitory activities of NAC to the purified NDM-5 were conducted. With the addition of set concentration of NAC, the hydrolytic activity of NDM-5 was remarkably decreased at the concentration of 4.8 mmol/L (**Fig. 5D**). The inhibitory activity was analyzed by detecting the various ODs of meropenem hydrolysis for NDM-5, and the IC_50_ was 5.55 mM (**Fig. 5E**). The potential competitive inhibitor, NAC, showed a K_i_ against NDM-5 of 2.89 mM (**Fig. 5F**).

### The accumulation of meropenem in bacteria promoted by NAC

The mechanism underlying the inhibition of biofilm formation and NDM activity have been elucidated. However, the important issue concerning the intracellular accumulation of meropenem remains unresolved. Meropenem, a β-lactam antibiotic that inhibits cell wall synthesis, requires sufficient intracellular concentration and other membrane variations for its efficacy against bacteria. First, the *N*-phenyl-1-naphthylamine fluorescent probe was utilized as a tool to assess the effect of NAC on outer membrane permeability. The results demonstrated that NAC increased the permeability of the outer membrane in a dose-dependent manner, with the concentraction of NAC (**Fig. 6A**). This suggests the potential for NAC to make the outer membrane more permeable, thereby, resulted in an increase of meropenem in the bacteria. Second, the DiSC_3_(5) fluorescent probe was used to analyze the influence of NAC on the cytoplasmic membrane. The result of the membrane depolarization assay indicated that NAC-dependent increases in fluorescence suggested a disruption of the electric potential (**Fig. 6B**). Third, the effects of NAC on cell membrane integrity, proton motive force, ATP detection and EtBr efflux pump were analyzed. NAC exposure resulted in a concentration-dependent significant reduction of intracellular ATP (**Fig. 6C**) and increased the inhibitory effect on Etbr efflux (**Fig. 6D**), respectively. The former may affect anabolism and catabolism, or inhibit the biofilm, while the latter may affect the accumulation of antibiotic. Conversely, there were no significant differences in the evaluation of NAC’s influence on cell membrane integrity and proton motive force by using propidium iodide and BCECF AM probes (**Fig. S1**), respectively. The direct evidence of meropenem accumulation, with a concentraction-dependent manner of NAC, was convincingly demonstrated by HPLC (**Fig. 6E**).

**Fig 6.**
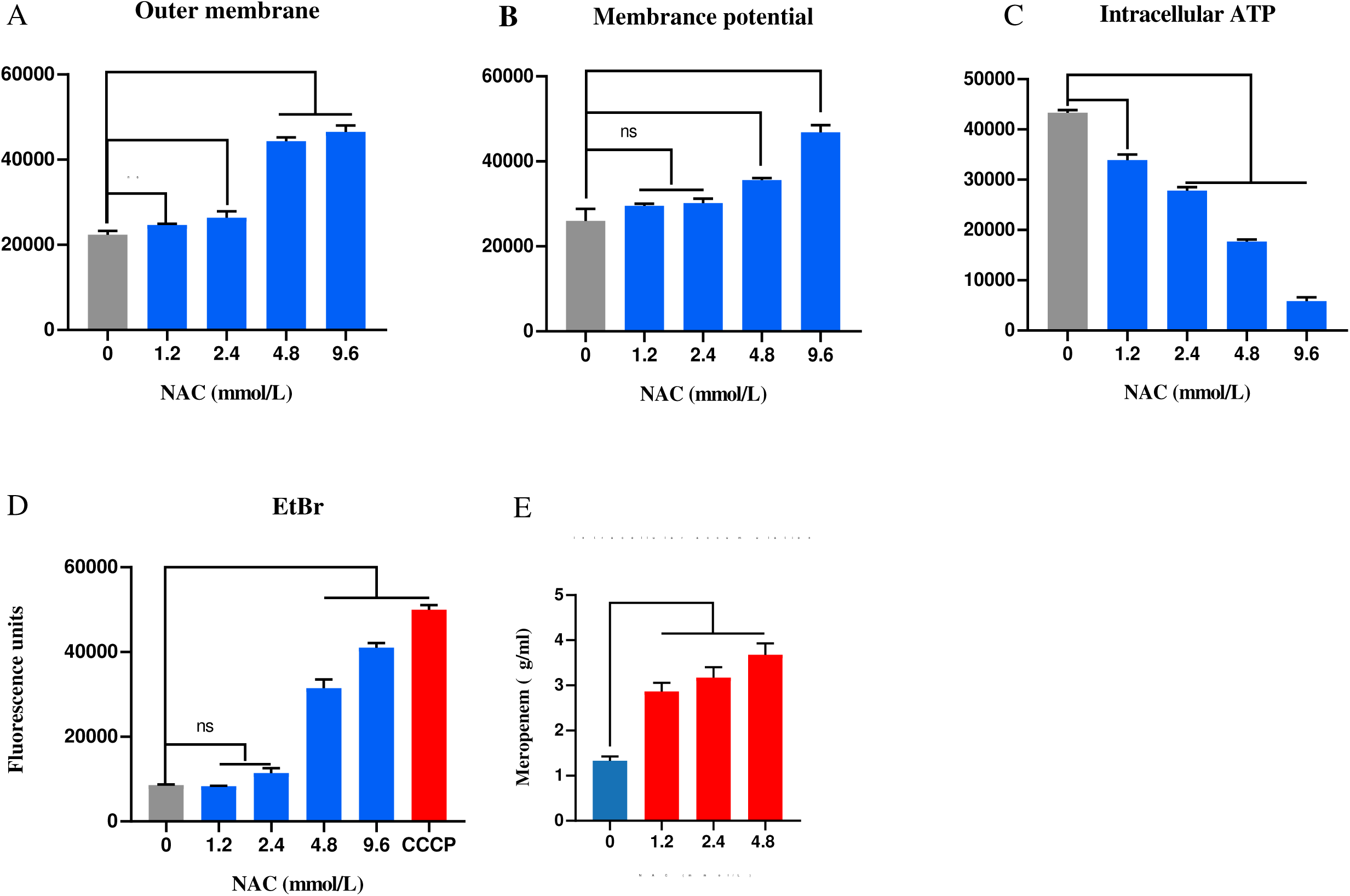
The potential mechanism of the meropenem and NAC combination. (A) NAC disrupts the bacterial outer membrane of *E. coli* 1911034 in a concentration-dependent manner. (B) NAC affects the membrane potential of *E. coli* 1911034 in a concentration-dependent manner. (3) Reduction of intracellular ATP treated with various concentrations of NAC. (D) Inhibition of the efflux pump treated with various concentrations of NAC. (E) Intracellular accumulation of meropenem by meropenem alone or plus NAC combination tests through HPLC. All data obtain from three biology replicates and the significances were evaluated by Prism 8 (GraphPad, USA) (**** P < 0.0001, *** P < 0.001 and ** P < 0.01).

### Conclusions

In Conclusions, Our results showed that the synergistic effect of the combination of meropenem and NAC could kill NDM-positive *E. coli*, and a specific concentration of NAC could restore the meropenem susceptibility *in vitro*. Further studies disclosed that it influences the biofilm, inhibits gene expression of *bla*_NDM-5_ and the hydrolytic activity of NDM-5 to meropenem as well as increases the intracellular accumulation of meropenem in a *bla*_NDM-5_ positive *E. coli* strain. These findings require further investigations for potential therapeutic applications of meropenem and NAC combinations. To the best of our knowledge, this is the first report of a synergistic activity of the combinations of meropenem and NAC in restoring meropenem susceptibility of *bla*_NDM-5_-carrying *E. coli*.

### Statistical analysis

Statistical analysis of data from the time–kill curve, antibiofilm assay and transcriptomic analysis, etc., was performed using a Dunnett’s multiple comparisons test on Prism 8 (GraphPad, USA), performed according to controls in each dataset. P < 0.0001, P < 0.01 and P < 0.05 were considered statistically significant in this study.

## Acknowledgements

This work was supported by a Special Fund for Young Talents in Henan Agricultural University (no.30601669), the China Postdoctoral Science Foundation (no. 2018M642751) and Jiangsu Agriculture Science and Technology Innovation Fund (CX(24)3078).

## Author contributions

Yuzhe Zhao, Data curation, Formal analysis, Investigation, Methodology, Validation,Writing-original draft. Xinyu Guo, Formal analysis, Investigation Methodology, Validation. Chenglong Li, Data curation, Formal analysis, Investigation, Visualization. Yixin Wang, Investigation; Methodology, Visualization. Lijuan Chang, Investigation, Visualization, Methodology. Yage Sun, Formal analysis, Validation, Investigation. Xiaoming Wang, Investigation, Formal analysis, Funding acquisition. Yanling Wang, Investigation, Formal analysis, Validation. Stefan Schwarz, Supervision, Project administration, Writing-review and editing. Dexi Li, Conceptualization, Data curation, Project administration, Funding acquisition, Supervision,Writing-original draft, Writing-review and editing. Xiang-Dang Du, Conceptualization, Supervision, Project administration; Writing-review and editing.

---

**Fig S1.**
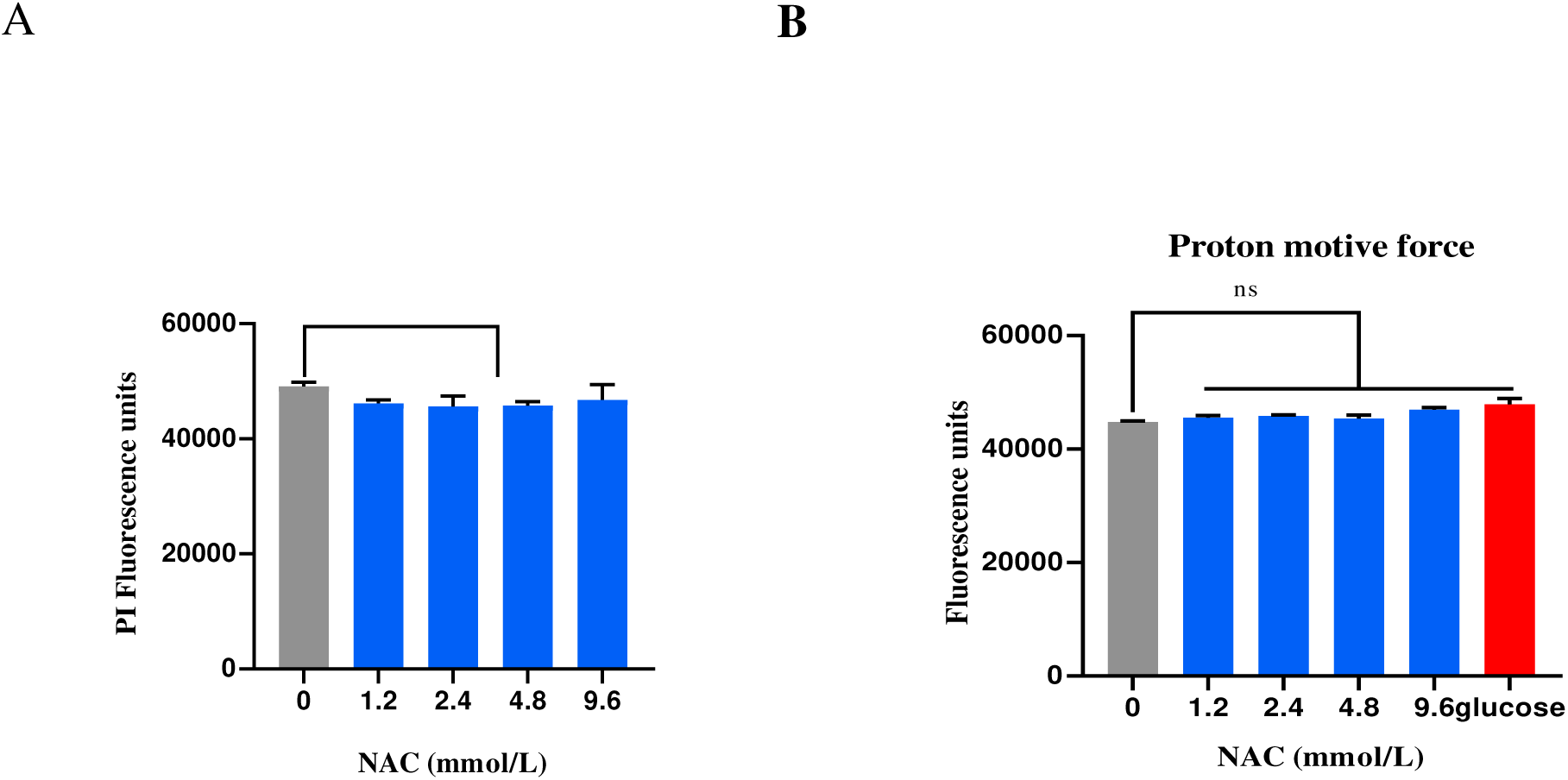
The potential mechanism by meropenem and NAC combination. (A) NAC has no effect on bacterial cytoplasmatic membrane after treatment with various concentrations of NAC in *E. coli* 1911034. (B) NAC has no effect on proton motive force of *E. coli* 1911034.

